# Hfq-assisted RsmA regulation is central to *Pseudomonas aeruginosa* biofilm polysaccharide PEL expression

**DOI:** 10.1101/434555

**Authors:** Yasuhiko Irie, Victoriia Murina, Vasili Hauryliuk, Victoria Shingler

## Abstract

Expression of biofilm-associated genes is controlled by multiple regulatory elements, allowing bacteria to appropriately switch between sessile and motile lifestyles. In *Pseudomonas aeruginosa,* the post-transcriptional regulator RsmA has been implicated in the control of various genes including those related to biofilms, but much of the evidence for these links is limited to transcriptomic and phenotypic studies. RsmA binds to target mRNAs to modulate translation by affecting ribosomal access and/or mRNA stability. Here we trace a global regulatory role of RsmA to the inhibition of Vfr – a transcription factor that controls a transcriptional regulator FleQ. FleQ directly controls biofilm-associated genes that encode the PEL polysaccharide biosynthesis machinery. Furthermore, we show that RsmA cannot bind *vfr* mRNA alone, but requires the RNA chaperone protein Hfq. This is the first example where a RsmA protein family member is demonstrated to require another protein for RNA binding.

## INTRODUCTION

The opportunistic pathogen *Pseudomonas aeruginosa* can be isolated from a wide range of environmental niches, in large part owing to its versatile metabolic capabilities. It is also proficient in colonizing various eukaryotic organisms and can cause both acute and chronic infections, with the latter often associated with biofilm-like modes of growth at the sites of infection (Gellatly and Hancock, 2013). *P. aeruginosa* is also highly competitive against other microbial species, with its adaptability and competitive fitness being aided by the production of various secondary metabolites and virulence factors (Tashiro et al., 2013;Khare and Tavazoie, 2015). Many of these cellular processes are inter-regulated by multiple regulatory pathways, presumably to provide appropriate response mechanisms to a wide range of environmental cues. One of the central regulators that determine *P. aeruginosa* behavior is the post-transcription factor RsmA (<u>R</u>egulator of <u>S</u>econdary <u>M</u>etabolite) as evidenced by its null mutant phenotypes linked to motility (Heurlier et al., 2004), virulence (Mulcahy et al., 2006;O’Grady et al., 2006;Mulcahy et al., 2008), biofilm (Mulcahy et al., 2006;Irie et al., 2010), growth (Irie et al., 2010), and secreted products (Pessi et al., 2001;Heurlier et al., 2004).

RsmA belongs to the RsmA/CsrA family of dimeric RNA-binding proteins (Gutiérrez et al., 2005;Rife et al., 2005;Schubert et al., 2007), homologues of which are found across a wide range of both Gram-negative and Gram-positive bacterial species (White et al., 1996): in some species including *E. coli*, they are called CsrA, while they are named RsmA in other species such as *P. aeruginosa*. RsmA/CsrA proteins inhibit translation of specific target mRNAs by binding to the ribosome binding sites, typically overlapping either the Shine-Dalgarno or the start codon (Romeo et al., 2013). However, in a few cases, they can also serve as positive regulators of translation by altering the secondary structures of bound RNAs (Patterson-Fortin et al., 2013;Ren et al., 2014) or protecting RNAs from degradation (Yakhnin et al., 2013).

Many RsmA/CsrA-targeted mRNAs are reported to have altered RNA decay rates. Several studies attempted to exploit this property to determine the global *P. aeruginosa* RsmA regulon using microarrays (Lawhon et al., 2003;Burrowes et al., 2006;Brencic and Lory, 2009). However, this approach cannot distinguish directly regulated genes from indirectly regulated gene sets that result from cascade regulation, nor can it identify target RNAs that do not exhibit changes in their stability (Baker et al., 2007;Pannuri et al., 2012). For example, in *P. aeruginosa*, direct RsmA-mediated translation inhibition has been demonstrated for the *P. aeruginosa*-specific biofilm polysaccharide operon *psl* without any effect on the level of transcription (Irie et al., 2010). Hence, the *psl* operon was not prominently featured in transcriptomic studies despite a strong biofilm phenotype (Burrowes et al., 2006;Brencic and Lory, 2009), raising concerns of what is currently considered to be ‘the RsmA regulon’ of *P. aeruginosa*. More direct global analyses of CsrA using techniques such as CLIP-Seq, RIP-Seq, and ribosome profiling have been performed in other organisms (Holmqvist et al., 2016;Potts et al., 2017;Sahr et al., 2017), but to our knowledge, such biochemical techniques to analyze the RsmA regulon have not been performed for *P. aeruginosa*.

Here, we present evidence that RsmA serves a master regulatory role through several intermediate transcription factors that are known to control different cellular processes. Through cascade regulation, originating from a direct effect on *vfr* mRNA, RsmA indirectly affects biofilm polysaccharide expression. Furthermore, we discovered an unexpected regulatory interplay between RsmA and the RNA chaperone protein Hfq on the target mRNA, whereby RsmA could only bind in the presence of Hfq. To our knowledge, this is the first example where RsmA/CsrA is unable to bind to its target alone. Because Hfq is known to facilitate RNA degradation (Bandyra and Luisi, 2013), these results may provide a mechanistic insight into RsmA-associated mRNA stability changes.

## RESULTS

### RsmA is a post-transcriptional regulator of *vfr* through which *fleQ* and *pel* transcripts are indirectly regulated

*P. aeruginosa* produces at least three different extracellular biofilm polysaccharides: alginate, PEL, and PSL (Ryder et al., 2007). PEL and PSL are co-regulated by several factors including the secondary messenger molecule c-di-GMP via the c-di-GMP-binding transcriptional regulator FleQ (Hickman and Harwood, 2008) and possibly also quorum sensing (Sakuragi and Kolter, 2007;Gilbert et al., 2009). Similar to PSL, PEL has long been considered to be regulated by RsmA (Gooderham and Hancock, 2009) based on a model originally proposed from a microarray study (Goodman et al., 2004). However, unlike the *psl* operon transcript, which was genetically and biochemically demonstrated to be directly regulated by RsmA (Irie et al., 2010), no direct evidence for the *pel* operon has been presented.

Consistent with a previous microarray study (Brencic and Lory, 2009), here using a chromosomal transcriptional reporter, we found that *pel* is up-regulated in the Δ*rsmA* mutant as compared to the wild type (WT) background (Fig. 1A). However, the PSL polysaccharides are also over-produced in a Δ*rsmA* strain (Irie et al., 2010) resulting in elevated intracellular levels of c-di-GMP due to PSL signaling (Irie et al., 2012). Given that the *pel* genes are transcriptionally up-regulated by c-di-GMP (Hickman and Harwood, 2008;Baraquet et al., 2012) and the Δ*rsmA* strain has elevated c-di-GMP levels, it is plausible that the up-regulation of *pel* could be due to increased c-di-GMP levels. To uncouple *pel* expression from elevated c-di-GMP, we measured *pel* transcriptional reporter activities when introduced into a low c-di-GMP Δ*rsmA* Δ*pel* Δ*psl* triple mutant background (Irie et al., 2012). In this strain *pel* is still up-regulated (Fig. 1A), indicating that the up-regulation is caused by lack of RsmA rather than changes in intracellular c-di-GMP concentration.

**Fig. 1.**
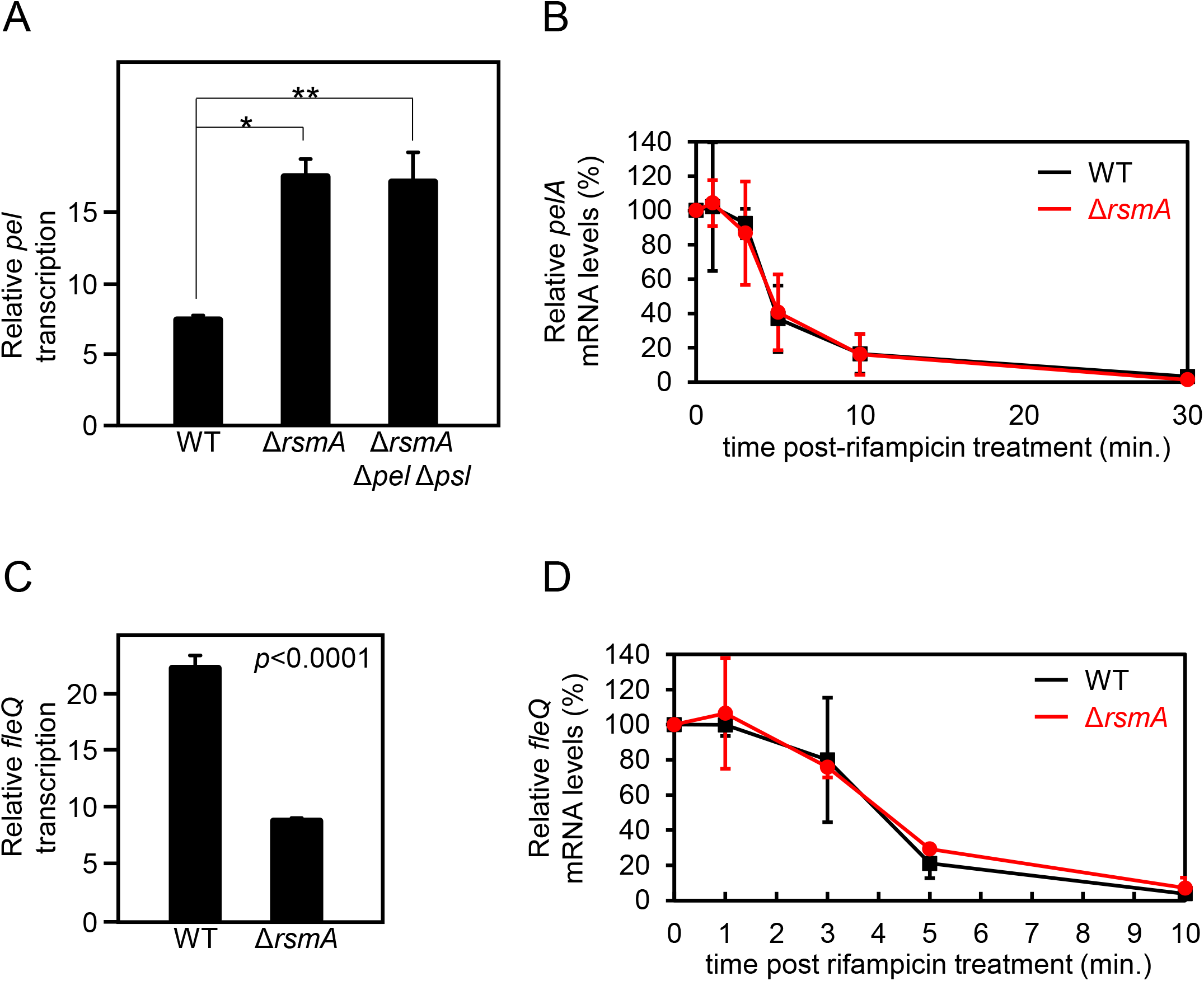
RsmA does not directly regulate *pel* nor *fleQ*. **A.** Single-copy transcriptional *lacZ* fusion constructs reveal up-regulation of *pel* transcripts in the Δ*rsmA* background as compared to wild type (WT). Δ*rsmA* strain overexpresses the PSL polysaccharides, which causes an elevation of intercellular concentration of c-di-GMP (Irie et al., 2012). Thus, we also tested a Δ*rsmA* Δ*pel* Δ*psl* strain, which has low c-di-GMP due to the absence of PSL (Irie et al., 2012). Note that *pel* is equally up-regulated in the Δ*rsmA* Δ*pel* Δ*psl* strain, demonstrating that the *pel* transcriptional phenotype of Δ*rsmA* is PEL-, PSL-, and c-di-GMP-independent \**p* = 0.0002; \*\**p* = 0.0013. **B.** Real-time quantitative PCR (qPCR) at different time points after rifampicin treatment show indistinguishable stabilities of the *pelA* transcript in the WT and Δ*rsmA* strains. C. Single-copy transcriptional *lacZ* fusion constructs reveal down-regulation of *fleQ* transcripts in the Δ*rsmA* background as compared to WT. D. qPCR at different time points after rifampicin treatment shows indistinguishable stabilities of the *fleQ* transcript in the WT and Δ*rsmA* strains.

Since direct regulation of mRNA levels by RsmA/CsrA often acts via altered mRNA stability (Romeo et al., 2013), we tested whether the apparent *pel* transcriptional changes between the WT and Δ*rsmA* strains is due to altered *pel* mRNA stability. This, however, is not the case (Fig. 1B), suggesting that the differences in *pel* expression are likely due to altered activity of the promoter. Because the RNA-binding protein RsmA is not a transcription factor, we tested the possibility of RsmA affecting expression of a known transcriptional regulator of *pel*, namely FleQ. As a well-characterized transcriptional activator of *pel* (Hickman and Harwood, 2008;Baraquet et al., 2012), potential RsmA-mediated alterations in expression of FleQ would be anticipated to affect transcription of the *pel* genes. Using a similar approach as described for *pel* above, we found that a transcriptional reporter fusion of *fleQ* gave lower levels in the absence of RsmA (Fig. 1C). This is consistent with previous data showing *pel* transcript levels are up-regulated in Δ*fleQ* backgrounds (Hickman and Harwood, 2008).

The data above could lend themselves to the interpretation that RsmA serves as a positive regulator of *fleQ*. While RsmA is more commonly known to be a translational repressor, this possibility is not without precedence. Examples of RsmA/CsrA proteins stimulating translation of target mRNAs include *phz2* and *moaA* in *P. aeruginosa* (Patterson-Fortin et al., 2013;Ren et al., 2014) and *flhDC* in *E. coli* (Yakhnin et al., 2013). While *P. aeruginosa* FleQ and *E. coli* FlhD4C2 share no sequence or mechanistic similarities, they are functional counterparts, with both being class I master regulators of flagellar biosynthesis in their respective organisms (Dasgupta et al., 2003). In addition, flagellar motility was previously found to be positively controlled by RsmA in *P. aeruginosa* (Heurlier et al., 2004), while in *E. coli,* CsrA has been shown to up-regulate flagellar gene expression through protecting *flhDC* mRNA from degradation (Yakhnin et al., 2013). It follows that if RsmA regulated *fleQ* in an analogous manner, *fleQ* mRNAs would be hyper-stabilized by RsmA and thus, less stable in the null mutant. However, no evidence of altered mRNA turnover rate was found between WT and Δ*rsmA* strains (Fig. 1D). Therefore, we hypothesized that the direct action of RsmA may lie even further upstream in a regulatory cascade that involved control of *fleQ* by Vfr.

Vfr acts as a transcriptional repressor of the *fleQ* promoter (Dasgupta et al., 2002). Therefore, to be consistent with the above phenotypes, RsmA would have to serve as a repressor of *vfr*. In line with this idea, comparison of *vfr* transcriptional reporter activities in WT and Δ*rsmA* strains revealed up-regulation of *vfr* transcripts in the absence of RsmA (Fig. 2A). To ensure that the regulatory cascade functioned as anticipated, we tested the prediction that over-expression of Vfr should ultimately result in increased activity of the *pel* transcriptional reporter. This is, indeed, the case (Fig. 2B). In contrast to *pel* (Fig. 1B) and *fleQ* (Fig. 1D), the transcript stability of *vfr* is significantly different between WT and Δ*rsmA* strains, such that *vfr* mRNAs are more stable in the absence of RsmA (Fig. 2C). These results indicate that RsmA affects *vfr* mRNA levels by mediating changes in turnover rates.

**Fig. 2.**
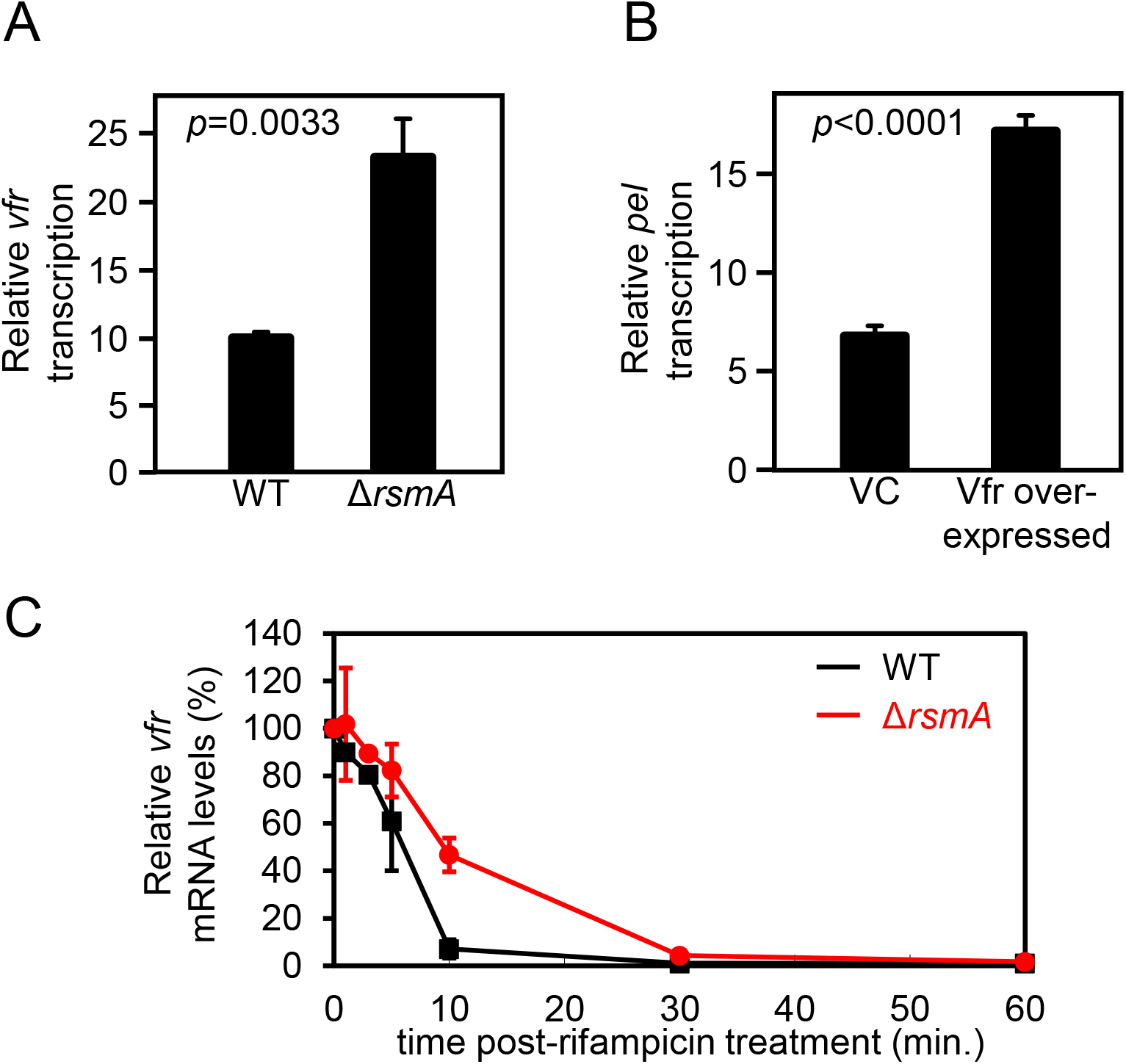
RsmA directly represses *vfr* by altering the mRNA stability. **A.** Single-copy transcriptional *lacZ* fusion constructs reveal up-regulation of *vfr* transcripts in the Δ*rsmA* background as compared to WT. **B.** The single-copy P_*pelA*_::*lacZ* transcriptional fusion has a higher activity in a Vfr over-expressing strain compared to the same strain harboring the vector control (VC). **C.** qPCR at different time points after rifampicin treatment show that *vfr* mRNA is less stable in WT as compared to the Δ*rsmA* strain.

### RsmA regulation of Vfr alters swimming and twitching motility

*P. aeruginosa* uses at least two major modes of motility: flagellar-driven swimming motility through liquid and surface ‘twitching’ motility using Type IV pilus (Burrows, 2012;Kazmierczak et al., 2015). Bacterial migration is an important aspect during *P. aeruginosa* biofilm development as initial attachment to a surface is thought to require flagella and mature microcolony formation has been attributed to Type IV pili functions (O’Toole and Kolter, 1998;Klausen et al., 2003). Thus, fine-tuned regulatory control of motility and biofilm genes is a necessity for effective *P. aeruginosa* adaptation, particularly when switching between motile and sessile lifestyles.

Because FleQ is a class I master regulator of flagellar genes (Dasgupta et al., 2003) and RsmA and Vfr lie upstream of FleQ in the regulatory cascade, lack of either protein should predictively impact flagellar motility, but in opposite ways. As would be predicted, Δ*rsmA* and Vfr over-expressing strains have reduced flagellar (swimming) motilities (Fig. 3A). Motilities of Δ*vfr* and RsmA over-expressing strain are comparable to the WT and vector control (VC) strains respectively probably because de-repression of flagellar genes do not necessarily lead to functional hyper-motility.

**Fig. 3.**
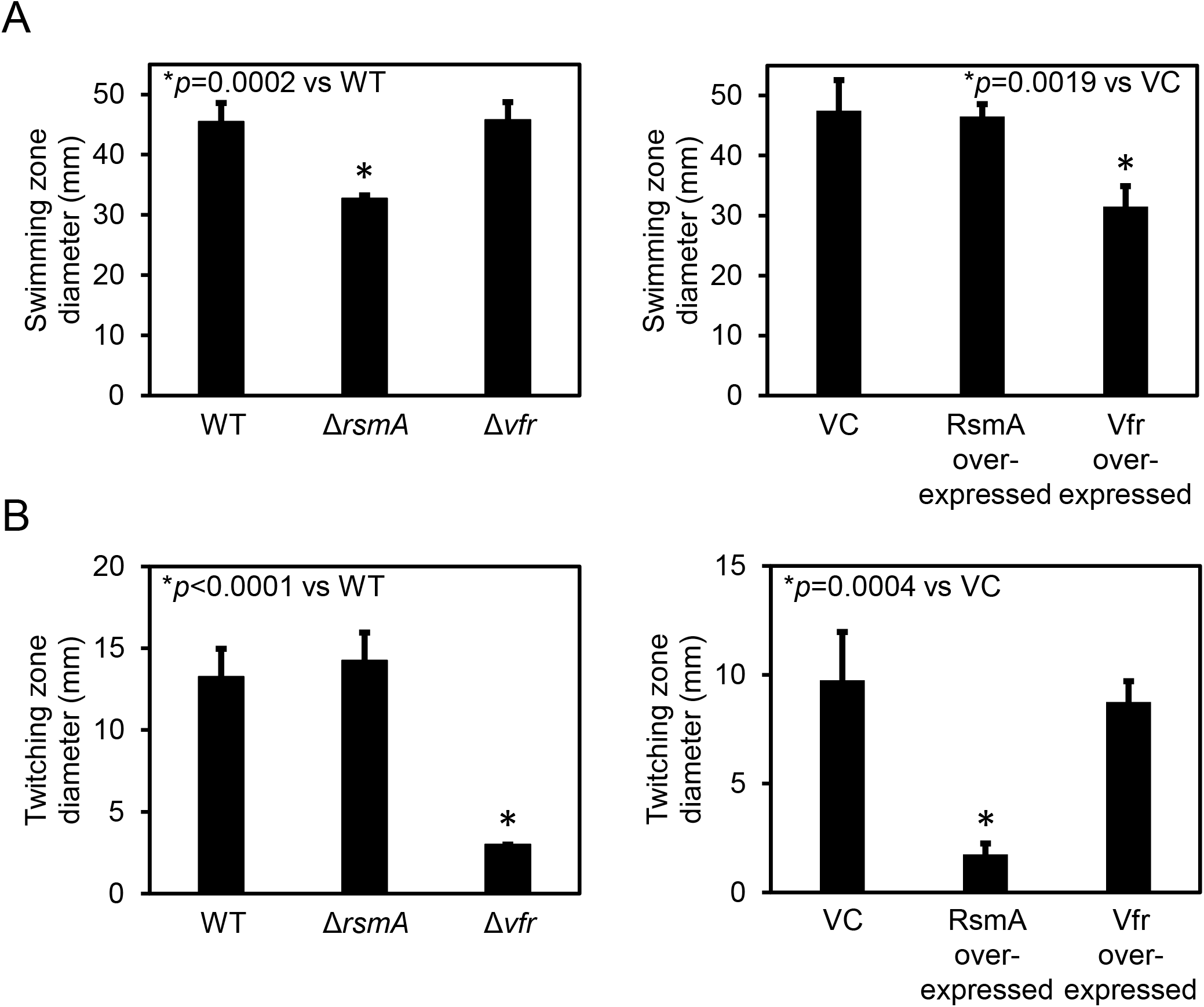
RsmA regulatory cascade indirectly affects flagellar and Type IV pilus-mediated motilities. **A.** Flagellar-mediated swimming motility is decreased for Δ*rsmA* and Vfr over-expressing strains. VC = vector control. Motility assays for VC, RsmA over-expressing strain, and Vfr over-expressing strains were performed in the presence of 300 μg/ml carbenicillin to ensure maintenance of the plasmids. **B.** Type IV pilus-mediated twitching motility is reduced for Δ*vfr* compared to WT. Δ*rsmA* motility is comparable to the WT strain probably because WT motility is maximized in this assay. Motility assays for VC, RsmA over-expressing strain, and Vfr over-expressing strains were performed in the presence of 300 μg/ml carbenicillin.

Mutants in *vfr* have previously been documented to be defective in Type IV pilus-dependent twitching motility (Whitchurch et al., 1996;Beatson et al., 2002) due to Vfr’s positive regulation of the AlgZR two-component system (Luo et al., 2015;Pritchett et al., 2015) required for expression of the Type IV pili genes (Lizewski et al., 2004;Belete et al., 2008). As shown in Fig. 3B, Δ*vfr* and RsmA over-expressing strains were both defective in twitching motility. Motilities of Δ*rsmA* and Vfr over-expressing strain are comparable to the WT and VC strains respectively probably because of similar reasons discussed with swimming motility above.

### Hfq is required for RsmA to bind to *vfr* mRNA

Given the evidence that RsmA may post-transcriptionally regulate *vfr*, we next examined the binding properties of RsmA to *vfr* mRNA. *P. aeruginosa* RsmA has a binding specificity for the CANGGAYG consensus sequence of its target mRNA (Schulmeyer et al., 2016), which is similar to the *E. coli* CsrA consensus sequences RUACARGGAUGU generated by SELEX (Dubey et al., 2005) and AUGGAUG generated by CLIP-Seq (Potts et al., 2017). The transcriptional start site for *vfr* lies 153 bases upstream of the translational start site (Fuchs et al., 2010). Inspection of the *vfr* leader sequence identified one putative RsmA-binding site overlapping the likely GGGA Shine-Dalgarno sequence (Fig. 4A).

**Fig. 4.**
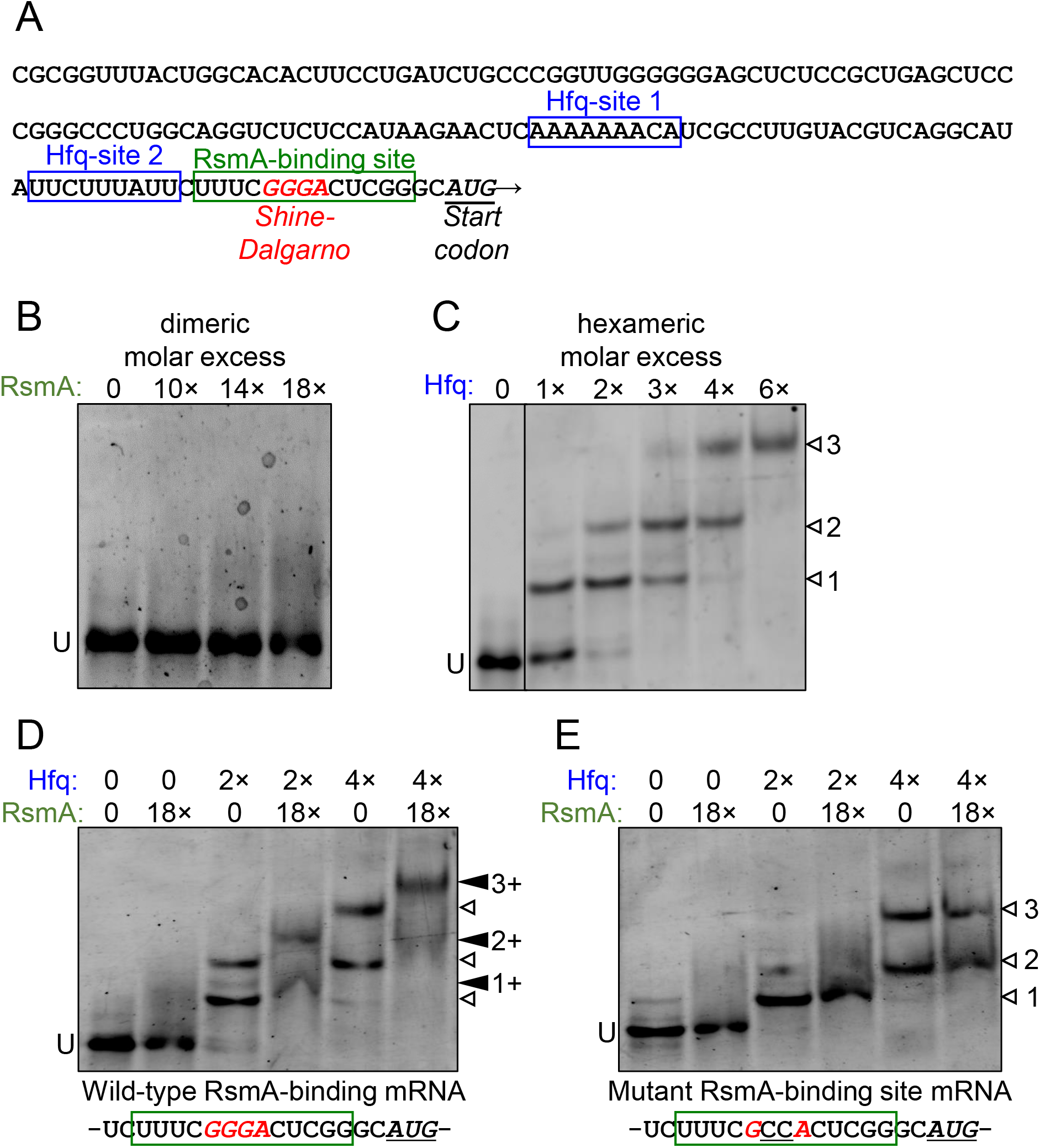
Direct targeting of *vfr* mRNA by RsmA requires Hfq. **A.** The leader sequence of the *vfr* transcript (Fuchs et al., 2010) possesses two putative Hfq-binding sites (blue boxes) and a RsmA-binding site (green box) overlapping the predicted Shine-Dalgarno sequence (red italics) upstream of the translational start site (underlined italics). **B.** RNA Electrophoretic mobility shift assay (EMSA) shows that RsmA does not bind to the *vfr* RNA when RsmA is present alone. Dimeric molar excess relative to RNA; U = unbound RNA. **C.** The *vfr* RNA produces three distinct band shifts (open arrow heads 1, 2, and 3) with increasing concentrations of Hfq. Hexameric molar excess relative to RNA. **D.** The presence of RsmA causes all Hfq-shifted bands to super-shift (filled arrow heads 1+, 2+, and 3+), indicating that RsmA binds to Hfq-bound RNA. Note that the RsmA concentration used (36× molar excess of RNA) is incapable of producing a shift when present alone (panel B). **E.** A GG→CC substitution within the RsmA-binding site abolishes RsmA super-shifts but does not alter Hfq-binding.

Purified RsmA protein directly binds *psl* mRNA (Irie et al., 2010) in RNA electrophoretic mobility shift assays (EMSA). However, no RsmA-binding was observed in similar assays with *vfr* RNA spanning the leader sequence (Fig. 4B). This result led us to consider the possibility that RsmA may require another factor to bind the *vfr* mRNA. RNA-binding chaperone protein Hfq was raised as a candidate for three reasons. First, the small RNA RsmY, which binds multiple RsmA proteins at high affinity to sequester and relieve RsmA-bound mRNAs (Kay et al., 2006), also has the capacity to be bound by Hfq (Sorger-Domenigg et al., 2007), although it was unclear from their study whether co-binding of both proteins occurred. Second, Hfq was recently identified to bind to *vfr* mRNA by global ChIP-Seq analyses (Kambara et al., 2018). Third, *in silico* analysis of the *vfr* leader sequence revealed two potential Hfq binding sites upstream of the putative RsmA-binding site (Fig. 4A).

Biochemical studies, primarily of *E. coli* Hfq, have shown that this hexameric protein complex has at least four regions which can all be involved in RNA-binding: the proximal face, the distal face, the rim, and the C-terminal tail (Updegrove et al., 2016). *P. aeruginosa* and *P. putida* Hfq have shortened C-termini compared to *E. coli* Hfq and may lack C-terminal tail-binding altogether (Milojevic et al., 2013). The proximal site preferentially binds to U-rich RNA sequence (Møller et al., 2002), the distal site binds to A-rich (ARN)n triplet repeats (Link et al., 2009), while the rim associates with UA-rich regions (Schu et al., 2015). The two potential Hfq-binding sites found within the *vfr* leader sequence (Fig. 4A) represent one A-rich ARN repeat region and one U-rich region.

RNA EMSA analyses show that *P. aeruginosa* Hfq binds *vfr* RNA, resulting in three distinct band shifts (Fig. 4C) indicative of two or more binding sites. Because wild-type *P. putida* Hfq showed identical binding patterns to *P. aeruginosa* Hfq on *vfr* RNA (Fig. S1A), we took advantage of previously characterized derivatives of *P. putida* Hfq, namely, a distal site mutant (HfqY25D; A-rich ARN-binding deficient) and a proximal site mutant (HfqK56A; U-rich-binding deficient) (Madhushani et al., 2015) to further analyze Hfq-binding to the *vfr* RNA. As seen in Fig. S1B, HfqY25D only recapitulates the second band shift while HfqK56A only recapitulated the first band shift. Notably, the third band shift is absent with both mutant derivatives of Hfq. We conclude that the first shift is caused by Hfq-binding to the U-rich region of the *vfr* leader sequence (Hfq-site 2 in Fig. 4A), the second shift is caused when the ARN repeats are bound by Hfq (Hfq-site 1 in Fig. 4A), while the third shift occurs when both sites are occupied simultaneously.

Having ascertained that Hfq does bind *vfr* RNA, we next added RsmA and Hfq simultaneously to the *vfr* RNA. As seen in Fig. 4D, this results in super-shifting of all three Hfq band shifts in the presence of RsmA, suggesting that binding of Hfq to either of its target sites is sufficient to allow RmsA-binding. Because the GG doublet within the consensus sequences for RsmA and CsrA binding are essential for their abilities to bind cognate target RNAs (Dubey et al., 2005;Irie et al., 2010), we made a CC substitution of the GG doublet within the predicted RsmA-binding region of *vfr* RNA. No RsmA-mediated super-shifting of Hfq band shifts was detected with the CC substituted RNA (Fig. 4E).

We infer from the findings above that Hfq-binding is a pre-requisite for RsmA to bind to its target within *vfr* mRNA. Together with the data in preceding sections, these results lead us to conclude: 1) that the direct binding of RsmA on *vfr* RNA requires the RNA chaperone protein Hfq and 2) that RsmA is an indirect regulator of biofilm polysaccharide locus *pel*, through its direct action on *vfr* propagated through a regulatory cascade from Vfr to FleQ.

## DISCUSSION

In this study, we report that *P. aeruginosa* RsmA requires the RNA chaperone Hfq to assist its binding to *vfr* mRNA to initiate a regulatory cascade that ultimately impacts *pel* biofilm polysaccharide genes. The requirement for Hfq was unexpected, since all other biochemical analyses of RsmA/CsrA family members document unaided binding to their target RNAs (Baker et al., 2002;Dubey et al., 2003;Wang et al., 2005;Lapouge et al., 2007;Irie et al., 2010;Patterson-Fortin et al., 2013;Yakhnin et al., 2013;Ren et al., 2014).

We identified two independent Hfq-binding sites upstream of the RsmA-binding motif, which overlaps the ribosome-binding site of *vfr* (Fig. 4A). Based on a mFOLD-predicted secondary structure (Zuker, 1989;2003), a large hairpin loop base-pairs the identified U-rich and A-rich Hfq-binding sites, resulting in a dsRNA structure that would predictively block both RsmA and the ribosome from accessing the mRNA (Fig. 5A). As depicted in Fig. 5B, the simplest model that we propose from our findings is that Hfq-binding is required to open the stem-loop to expose the RsmA-binding site. Subsequent binding of RsmA would directly block the Shine-Dalgarno sequence, preventing ribosome access and thereby inhibiting translation. We also provide evidence that *vfr* mRNA is hyper-stable in the absence of RsmA (Fig. 2C), indicating that Hfq- and RsmA-bound *vfr* mRNA may undergo more rapid degradation. It was previously suggested that Hfq may be able to directly recruit RNase E for the degradation of bound RNA in *E. coli* (Morita et al., 2005), and it is possible that a similar mechanism causes *vfr* mRNA instability.

**Fig. 5.**
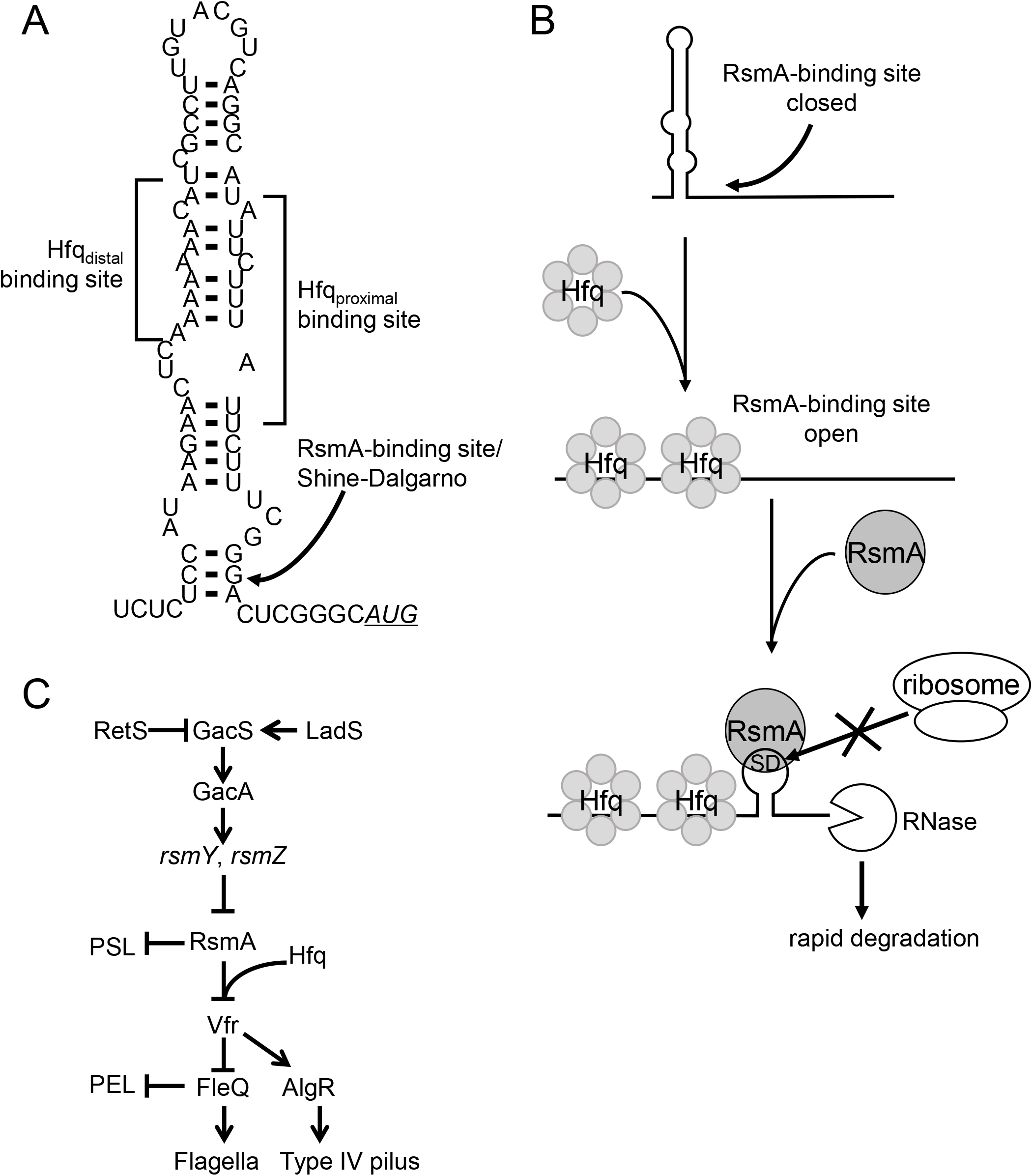
Model for Hfq-assisted RsmA regulation of *vfr* and its consequences. **A.** RNA secondary structure of the leader sequence region of *vfr* as predicted by mFOLD (Zuker, 1989;2003). The locations of *in silico*-detected Hfq- and RsmA-binding sites are indicated. Note that the RsmA-binding site located at the base of the stem loop structure overlaps with the Shine-Dalgarno sequence of the ribosome binding site. The AUG start codon is shown underlined in italics. **B.** Schematic illustration of a model for Hfq-assisted RsmA regulation of *vfr*. Upon Hfq-binding of *vfr* RNA, the stem loop structure shown in panel A is disrupted, exposing the RsmA-binding site. RsmA-binding then directly blocks the Shine Dalgarno (SD) sequence. RsmA-binding is depicted as producing a smaller stem-loop structure (Gutiérrez et al., 2005;Rife et al., 2005;Schubert et al., 2007), preventing ribosome access and thereby inhibiting translation. *In vivo*, the Hfq- and RsmA-bound mRNAs are presumed to be subsequently processed for degradation. **C.** Simplified overview of the Gac/Rsm pathway that controls RsmA levels and pertinent parts of its downstream regulon.

Due to its histidine-rich C-terminus, *E. coli* Hfq is a common contaminant of His-tagged proteins purified by nickel affinity chromatography after over-expression in *E. coli*, and visually undetectable (on SDS-PAGE) Hfq levels can critically influence *in vitro* analyses of RNA-binding (Milojevic et al., 2013;Moreno et al., 2015). Because the proteins used in this study were purified from over-expressing them in either *P. aeruginosa* (RsmA) or a Hfq null strain of *E. coli* (Hfq proteins), our analyses were not complicated by this issue. However, a large majority of analyses performed in past publications use C-terminal His-tagged RsmA/CsrA proteins derived from over-expression in *E. coli,* raising the possibility of *E. coli* native Hfq contamination. Given that Hfq binds *vfr* RNA with apparent high affinity (Figs. 4C and S1), it is plausible that Hfq-assisted RsmA-binding may be prevalent. Although such a possibility would not alter major conclusions and remains to be experimentally verified, it reveals provocative observations that comparison of transcriptomic studies done for RsmA, Hfq, and Vfr regulons in *P. aeruginosa* identify numerous overlaps of genes that were differentially expressed between WT and the respective null mutants (Wolfgang et al., 2003;Goodman et al., 2004;Burrowes et al., 2006;Brencic and Lory, 2009;Sonnleitner et al., 2018). Overlaps were also seen in *E. coli* between the Hfq and CsrA regulons (Potts et al., 2017), suggesting that interplays between the two RNA-binding regulators is not unique to *P. aeruginosa*.

A diverse range of phenotypes have been associated with RsmA functions in *P. aeruginosa* (Vakulskas et al., 2015), but phenotypic observations do not distinguish direct regulatory activities from indirect effects through regulatory cascades. Our identification of Hfq and RsmA action on *vfr* mRNA resulted from an initial aim to determine whether RsmA directly targeted *pel* RNA, as is the case for *psl* RNA (Irie et al., 2010). Our backtracking to trace the ultimate cause of RsmA effects on *pel* expression revealed a regulatory pathway originating from Vfr to the transcriptional regulator FleQ and from there to *pel* (Fig. 5C).

This work places the global post-transcriptional regulators RsmA and Hfq as a hub at the center of upstream and downstream regulatory pathways and thus provides a conceptually new framework to evaluate and interpret past work. The two component GacS/GacA system lies at the top of the hierarchy that controls RsmA-mediated regulation. Although the environmental cue(s) for activation is still unknown, GacS activity is modulated through interactions with RetS and LadS (Goodman et al., 2009;Kong et al., 2013;Chambonnier et al., 2016). GacA, in turn controls the production of the small RNAs – *rsmY* and *rsmZ* – which possess multiple high affinity binding sites to titrate RsmA away from target mRNAs (Heurlier et al., 2004;Sonnleitner et al., 2006). In light of a report indicating that Hfq also binds to *rsmY* (Sorger-Domenigg et al., 2007), it appears plausible that these small RNAs may also require Hfq to assist their functions.

In addition to biofilm-associated phenotypes, there are numerous other processes associated with RsmA in *P. aeruginosa*, including motility (Fig. 3) and acute virulence (Mulcahy et al., 2006;Intile et al., 2014). Many of these ‘RsmA-regulated factors and phenotypes’ are likely to be indirectly regulated through Vfr- and/or FleQ-initiated regulatory cascades. Based on our findings, it will be crucial to determine and distinguish direct versus indirect regulatory routes to gain a greater understanding of the RsmA regulon.

## MATERIALS AND METHODS

### Bacterial strains and growth conditions

Table S1 lists the bacterial strains used in this study. *E. coli* and *P. aeruginosa* strains were grown in lysogeny broth (LB) Lennox composition at 37°C unless specified otherwise. VBMM citrate medium (Hoang and Schweizer, 1997) was used for selecting *P. aeruginosa* post-conjugation. For *E. coli,* the following antibiotics concentrations were used: 50 μg·ml^-^ carbenicillin, 10 μg·ml^−1^ gentamicin, and 10 μg·ml^−1^ tetracycline. For *P. aeruginosa* strains: 300 μg·ml^−1^ carbenicillin, 100 μg·ml^−1^ gentamicin, and 100 μg·ml^−1^ tetracycline were used. Sucrose counter-selection for plasmids carrying the *sacB* gene used in *P. aeruginosa* strain constructions was performed by streaking colonies on LB agar (no salt) supplemented with 10% w/v sucrose. Plates were incubated at 30°C for 24 hours, after which the counter-selected colonies were confirmed for the loss of antibiotic resistance and mutations confirmed by PCR for double-cross-over genomic mutants.

### Strain constructions

Transcriptional fusion constructs were generated in single copy on the chromosome of *P. aeruginosa* strains via integration into the chromosomal *attB* site as previously published (Irie et al., 2010). In brief, promoter regions were PCR amplified using oligonucleotides in Table S2. PCR products were ligated between the EcoRI and BamHI sites of mini-CTX *lacZ* (Fig. S2). These plasmids were then introduced into *P. aeruginosa* by conjugation. After double site recombination, plasmid backbones were removed by FLP recombinase, and then strains cured of the pFLP2 plasmid by sucrose counter-selection.

### β-galactosidase assays

Quantitative β-galactosidase activities were assayed using Galacto-Light Plus kit (Thermo-Fisher) as previously published (Lequette et al., 2006). All *P. aeruginosa* cultures were grown in VBMM citrate at 37°C to exponential phase, and lysed using chloroform as previously described (Irie et al., 2010). β-galactosidase activity units were normalized to total proteins per ml as determined using Bradford assay reagents (Bio-Rad). Assays were performed in biological triplicates.

### Quantitative real-time PCR and RNA stability analyses

Real-time PCR was performed as previously described (Irie et al., 2012), using the oligonucleotides listed in Table S2. For RNA stability experiments, exponential phase *P. aeruginosa* cultured in VBMM citrate at 37°C were treated with 200 μg·ml^−1^ rifampicin (Lory, 1986). RNAs were extracted from 1 ml of the cultures at various time points as previously described (Irie et al., 2010) after the addition of rifampicin. RNA extractions were performed using the RNeasy Mini Kit (Qiagen) after treating the harvested cells with RNAprotect (Qiagen). Genomic DNA was removed using DNase I (Promega) and removal confirmed by PCR using primers designed against the *rplU* gene (Table S2). SuperScript III First-Strand Synthesis (Invitrogen) was used to synthesize cDNA as per manufacturer’s protocol using random hexamers. Quantitative real-time PCR was performed using SYBR Green PCR Master Mix (Thermo/Applied Biosystems). The internal control gene used was *ampR*. All experiments were done in biological quadruplicates.

### Motility assays

Motility assays were performed essentially as previously described (Shrout et al., 2006). For swimming motility, LB-Lennox plates containing 0.3% Bacto agar were inoculated with overnight cultures with a sterile inoculation needle, ensuring the needle tip was inserted approximately halfway into the agar but not to the plastic petri dish bottom, and incubated for 24 hours at 30°C. Swim ring diameters were measured for quantitation. For twitching motility, LB-Lennox plates containing 0.5% Bacto agar were inoculated by inserting a sterile inoculation needle through the agar until it touched the plastic bottom. Plates were incubated for 72 hours at 30°C. Subsequently, the agar was peeled off, and the plates were stained with 1% w/v crystal violet to visualize the twitching zone diameters prior to measurement (Déziel et al., 2001). All motility experiments were performed in at least biological quadruplicates each in technical quadruplicates.

### Protein expression and purification

Proteins were purified by previously established protocols. Briefly, over-expressed RsmA-His_6_ was purified from *P. aeruginosa* PAO1 bearing pUCP18::*rsmA*-His_6_ (Irie et al., 2010), while all Hfq-His_6_ proteins were purified from a Hfq null derivative of *E. coli* BL21(DE3) (Madhushani et al., 2015) carrying PT7 expression plasmids (pVI2344-2346 or pVI2357).

RsmA and Hfq over-expressing strains were grown in 2× YT broth to stationary or exponential phases, respectively. Cell pellets were re-suspended in lysis buffer (buffer A: 0.3 M NaCl, 20 mM Tris-HCl pH 8, 5 mM imidazole) supplemented with 1 tablet/L cOmplete^™^ EDTA-free protease inhibitor tablet (Sigma). Cells were lysed by Stansted Fluid Power SPCH ultra high-pressure cell disrupter/homogenizer. Cell debris were removed by centrifugation (35,000 rpm, 40 minutes, 4°C) and the supernatants loaded onto 1 mL HisTRAP HP (GE Healthcare) columns equilibrated in lysis buffer (buffer A). The column was washed with high salt buffer (buffer B: 5 M NaCl, 20 mM Tris-HCl pH 8, 5 mM imidazole) to remove RNA contamination and the proteins subsequently eluted with the linear gradient of imidazole (25 mM to 750 mM) in buffer C (buffer C: 0.5 M NaCl, 20 mM Tris-HCl pH 8). Fractions containing the desired proteins were pooled. For RsmA-His_6_, an additional ion exchange step using a 5 ml HiTrap SP HP column (GE Healthcare) equilibrated with buffer D (0.1 M NaCl, 20 mM Tris-HCl pH 8) was performed, with RmsA subsequently being eluted with 1 M NaCl.

All proteins were dialyzed across Slide-a-lyzer 10,000 MWCO cassettes (Thermo Fisher) against 2 lots of 2 L dialysis buffer (40 mM Tris-HCl pH 8, 600 mM NaCl, 2 mM EDTA) at 4°C for 2 days prior concentration using 10 MWCO Centricons (Amicon). Protein concentrations were determined using the BCA Protein Assay Kit (Pierce/Fisher) and purity confirmed by SDS-PAGE. Protein were stored at −20°C in final storage buffer (20 mM Tris-HCl pH 8, 300 mM NaCl, 1 mM EDTA, 40% glycerol) until use.

### RNA synthesis

Wild-type and mutant *vfr* mRNAs were generated using Ambion’s MEGAscript kit as recommended for *in vitro* transcription from the PT7 promoter of linearized plasmids. Reactions (total 40 μl) containing 1 μg of EcoRV-linearized pVI2358 or pVI2359, were incubated for 6 hours prior to 15 minutes DNase I treatment. Reactions were terminated by adding 230 μl RNase-free H2O and 30 μl stop solution (5 M ammonium acetate; 100 mM EDTA). The resulting 233 nt RNAs (encompassing co-ordinates −106 to +116 relative to the A of the initiation codon of *vfr*) were extracted twice with phenol:chloroform:IAA (25:24:1) and once with chloroform prior to precipitation and resuspension in 50 μl RNase-free H_2_O.

### RNA electrophoretic mobility shift assays (EMSAs)

Reactions (total 10 μl) contained 32 nM *in vitro* transcribed RNA with the indicated concentrations of RsmA and/or Hfq. RsmA and Hfq molarities given here are for the dimeric and hexameric complexes respectively. RNA was first heated at 80°C for 5 min and then immediately chilled in an ice-water bath. Additional final reaction mixture components were as follows: 10 mM HEPES pH 7.9, 2 mM MgCl_2_, 90 ng yeast total RNA (Fisher), 4U RNasin, and 35 mM KCl. Binding reactions were performed at 20°C for 40 min prior to adding 2.5 μl loading buffer (40% sucrose). Reactions were placed on ice and analyzed on 5% TBE Criterion pre-cast gels (Bio-Rad) with electrophoresis at 100V for ≈130 min at 4°C. Gels were stained at room temperature with 1:10,000 TBE-diluted SYBR Gold (Fisher) in a light-protected container with agitation. Images were captured using Typhoon FLA 9500 (GE Healthcare).

### Statistics

Presented data are the mean with standard deviations of the data collected. Student’s *t*-test using the GraphPad Prism software was used to calculate *p*-values.

## Supporting information

Supplemental Table 1, 2, supplemental figure legends, and supplmental material reference list

## ACKNOWLEDGEMENTS

Plasmid pMMBGW:RBS-*vfr* was a kind gift from Joe J. Harrison. YI would like to thank Matt Parsek for his more than a decade of scientific and moral support that led to the culmination of this project. The manuscript partially overlaps with a preprint at BioRxiv (Irie Y *et al.*, 2019).

## AUTHOR CONTRIBUTIONS

YI and VS designed and performed the experiments and analyzed the data; VM and VH assisted in purification of the RsmA and Hfq proteins. YI and VS wrote the manuscript with input from VH and VM.

---

**Fig. S1.**
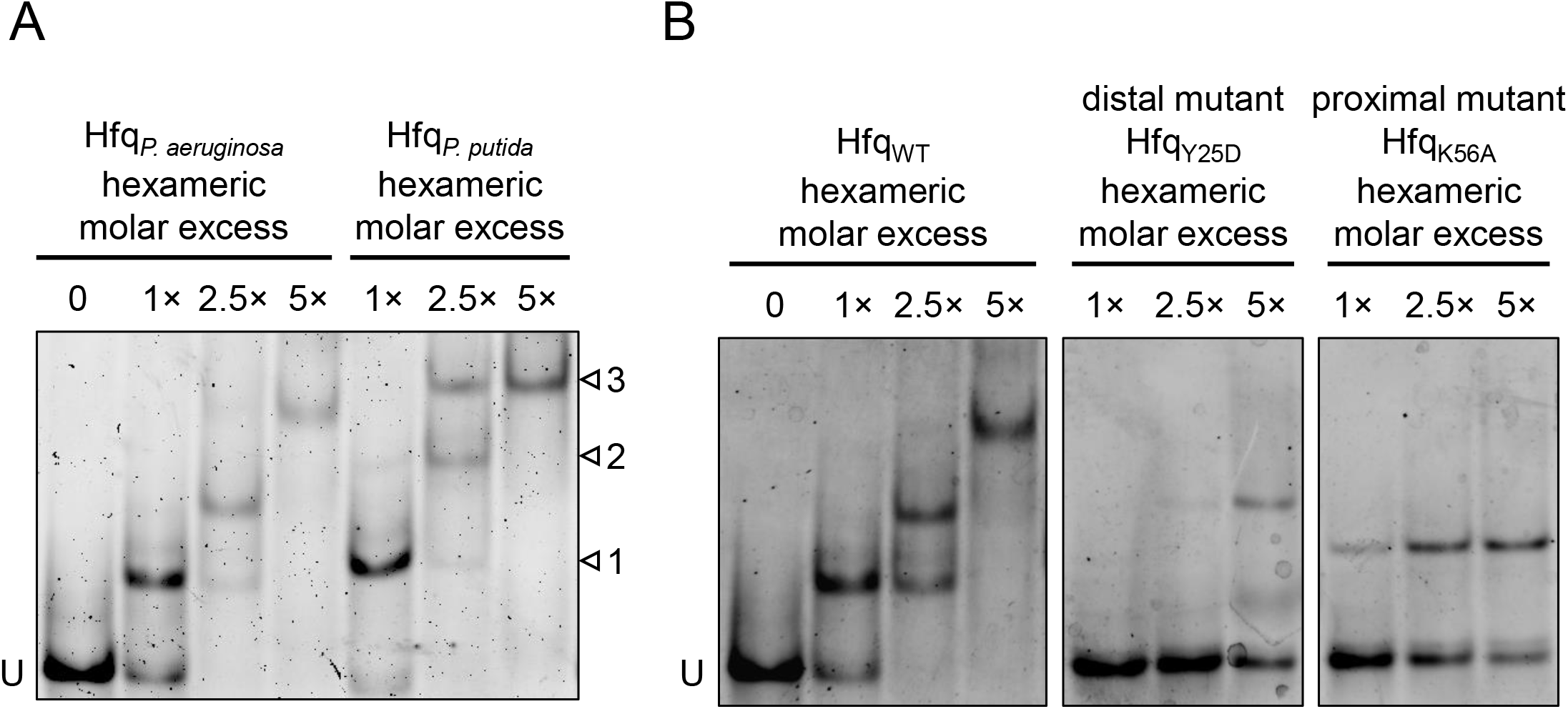

**Fig. S2.**
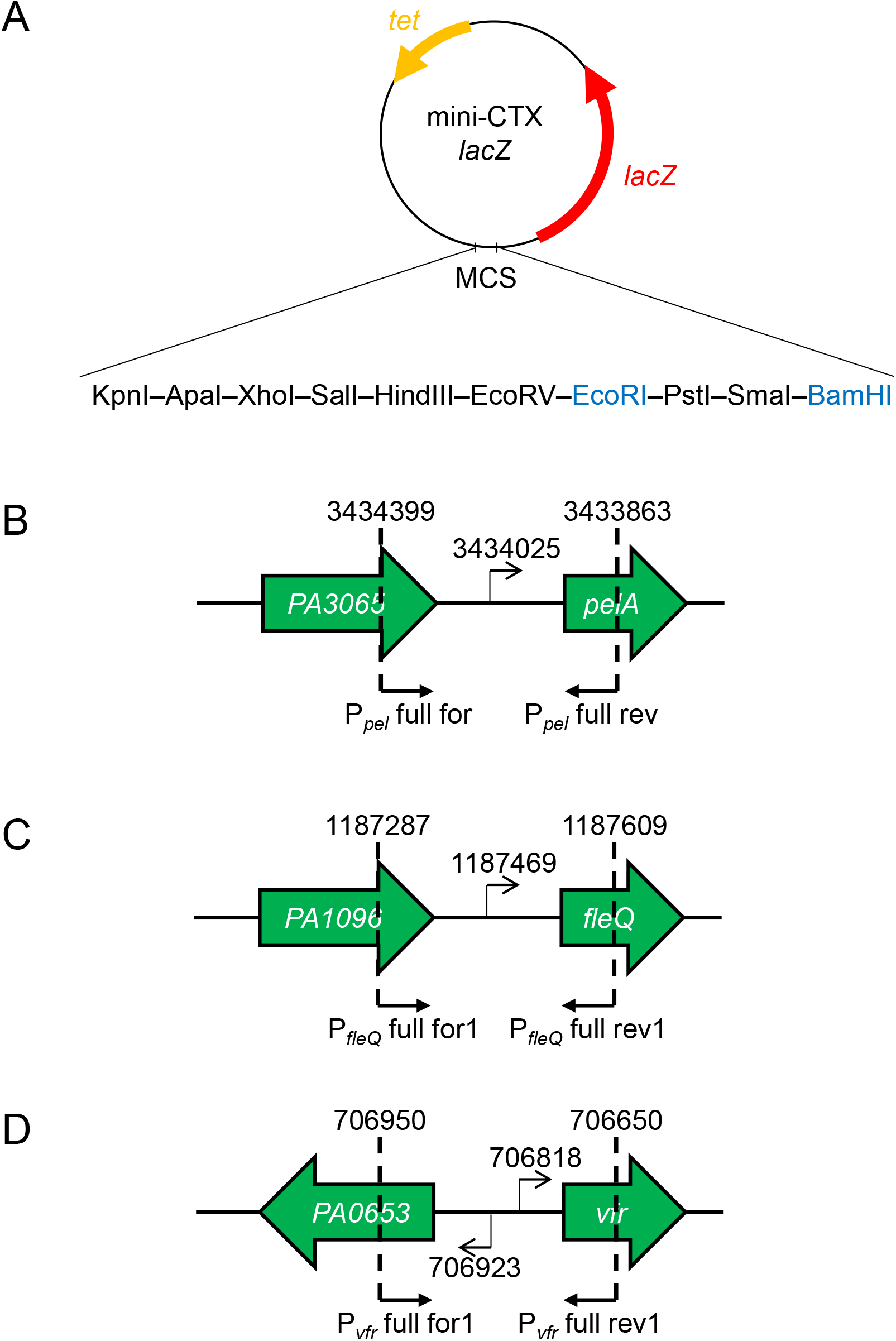

