## Supplemental Table 1, 2, supplemental figure legends, and supplmental material reference list for "Hfq-assisted RsmA regulation is central to *Pseudomonas aeruginosa* biofilm polysaccharide PEL expression"

SUPPLEMENTARY MATERIALS

SUPPLEMENTARY TABLES

Table S1

Bacterial strains and plasmids used in this study.

| Strains/plasmids | Description | Source/reference |
|---|---|---|
| *Pseudomonas aeruginosa* | | |
| PAO1 | wild type | (1) |
| Δ*rsmA* | Δ*rsmA*$_{-57\rightarrow73}$::*FRT* | (2) |
| Δ*rsmA* Δ*pel* Δ*psl* | Triple mutant | (2) |
| Δ*vfr* | In-frame *vfr* deletion | (3) |
| PAO1 $P_{pelA}$ full::*lacZ* trx | *lacZ* transcriptional fusion construct | this study |
| Δ*rsmA* $P_{pelA}$ full::*lacZ* trx | *lacZ* transcriptional fusion construct | this study |
| Δ*rsmA* Δ*pel* Δ*psl* $P_{pelA}$ full::*lacZ* trx | *lacZ* transcriptional fusion construct | this study |
| PAO1 $P_{fleQ}$ full::*lacZ* trx | *lacZ* transcriptional fusion construct | this study |
| Δ*rsmA* $P_{fleQ}$ full::*lacZ* trx | *lacZ* transcriptional fusion construct | this study |
| *Escherichia coli* | | |
| BL21(DE3) -Δ*hfq*::*cat* S1 | Hfq null T7 polymerase expression host | (4) |
| DH5 | cloning strain<br>*endA1 hsdR17*($r_k^-m_k^+$) *supE44 thi-1 recA1 gyrA96 relA1* (φ80*dlac*Δ*(lacZ)M15*) | (5) |
| NEB5α | cloning strain<br>*fhuA2* Δ*(argF-lacZ)U169 phoA glnV44* φ*80*Δ(*lacZ*)M15 *gyrA96 recA1 relA1 endA1 thi-1 hsdR17* | New England BioLabs |
| S17-1 λ*pir* | conjugation donor<br>*recA pro hsdR* RP4-2-Tc::Mu-Km::Tn7 λ*pir* | (6) |
| DB3.1 | conjugation donor<br>*thi thr leu tonA lacY supE recA*::RP4-2-Tc::Mu $Km^R$ | (7) |

Plasmids

| | | |
|---|---|---|
| pMMBGW | Gateway adapted pMMB67EH<br>Amp/$Carb^R$ | (8, 9) |

| | | |
|---|---|---|
| pMMBGW::RBS-*vfr* | *vfr* over-expression construct Amp/Carb$^R$ | this study |
| pUC57 | Cloning vector; Amp/Carb$^R$ | GenScript |
| pVI2358 | pUC57-based plasmid for $P_{T7}$ expression of the *vfr* RNA leader sequence. Insert was custom synthesized (GenScript) as an EcoRI/EcoRV fragment, encompassing the T7 promoter sequence and *vfr,* co-ordinates -106 to +116 relative to the A of the initiation codon. Carb$^R$ | this study |
| pVI2359 | As above but with a GG at -9 to -10 exchanged to CC within the *vfr* leader sequence Carb$^R$ | this study |
| pFLP2 | FLP recombinase expressing plasmid Amp/Carb$^R$ | (10) |
| pUCP18 | *P. aeruginosa - E. coli* shuttle vector Amp/Carb$^R$ | (11) |
| pRsmA ox | RsmA over-expression plasmid (pUCP18 backbone) Amp/Carb$^R$ | (2) |
| pUCP18::*rsmA*-His$_6$ | RsmA-His$_6$ over-expression plasmid Amp/Carb$^R$ | (2) |
| mini-CTX *lacZ* | *lacZ* transcriptional fusion *attB* integration construction plasmid Tet$^R$ | (12) |
| pET3H | T7 promoter expression vector; Amp/Carb$^R$ | (13) |
| pVI2344 | $P_{T7}$-PP *hfq*-His pET3H-based expression plasmid; Amp/Carb$^R$ | (4) |
| pVI2345 | $P_{T7}$-PP *hfq-Y25D*-His pET3H-based expression plasmid; Amp/Carb$^R$ | (4) |
| pVI2346 | $P_{T7}$-PP *hfq-K56A*-His pET3H-based expression plasmid; Amp/Carb$^R$ | (4) |
| pVI2357 | $P_{T7}$-PA *hfq*-His pET3H-based expression plasmid; Amp/Carb$^R$ | this study |

Amp = Ampicillin; Carb = Carbenicillin; Km = Kanamycin; Tet = Tetracycline

Table S2

Oligonucleotide primers used in this study. Engineered restriction sites are underlined. Genome co-ordinates are based on the annotation of PAO1 on www.pseudomonas.com (14, 15).

| Oligonucleotide primer | Used for | Genome co-ordinates | Sequence (5' to 3') |
|---|---|---|---|
| $P_{pel}$ full for | $P_{pel}$ full::*lacZ* construction | 3434399-3434376 | CGGCGAATTCCTGGTGCGGTTCCTCGCACGCAAC |
| $P_{pel}$ full rev | $P_{pel}$ full::*lacZ* construction | 3433863-3433883 | GATCGGATCCACGGCGATTCCTTTCTTGCTG |
| $P_{fleQ}$ full for1 | $P_{fleQ}$ full::*lacZ* construction | 1187287-1187306 | CGGCGAATTCCTACCAGATGTTCGGATAAG |
| $P_{fleQ}$ full rev1 | $P_{fleQ}$ full::*lacZ* construction | 1187609-1187589 | GATCGGATCCAAGAGTTTGGTTTCGCGCCAC |
| $P_{vfr}$ full for1 | $P_{vfr}$ full::*lacZ* construction | 706950-706934 | CGGCGAATTCCTTTCATCGTTCAGACT |
| $P_{vfr}$ full rev1 | $P_{vfr}$ full::*lacZ* construction | 706650-706672 | GATCGGATCCGGTGTGTGGGTAATAGCTACCAT |
| *rplU* forward | Checking for gDNA contamination in RNA prep | 5116619-5116600 | CGCAGTGATTGTTACCGGTG |
| *rplU* reverse | Checking for gDNA contamination in RNA prep | 5116315-5116334 | AGGCCTGAATGCCGGTGATC |
| *ampR*-F | RT-PCR | 4593527-4593511 | GCGCCATCCCTTCATCG |
| *ampR*-R | RT-PCR | 4593473-4593491 | GATGTCGACGCGGTTGTTG |
| *pelA*-F | RT-PCR | 3432090-3432070 | CCTTCAGCCATCCGTTCTTCT |
| *pelA*-R | RT-PCR | 3431973-3431992 | TCGCGTACGAAGTCGACCTT |
| *fleQ* RT for1 | RT-PCR | 1187644-1187666 | CTGGCAGTCATTCTCAACTTCCT |
| *fleQ* RT rev1 | RT-PCR | 1187706-1187689 | TCGCCAATCCTCGCTGTT |
| *vfr* RT for1 | RT-PCR | 706504-706489 | GACGGCCGCGAAATGA |
| *vfr* RT rev1 | RT-PCR | 706446-706462 | CCCAGCTCGCCGAAGAA |

SUPPLEMENTARY FIGURE LEGENDS

**Fig. S1 RsmA indirectly down-regulates *pel* transcript levels.**

A. Single-copy transcriptional *lacZ* fusion constructs reveal up-regulation of *pel* transcripts in the Δ*rsmA* background as compared to wild type (WT). Δ*rsmA* strain overexpresses the PSL polysaccharides, which causes an elevation of intercellular concentration of c-di-GMP (16). Thus, we also tested a Δ*rsmA* Δ*pel* Δ*psl* strain, which has low c-di-GMP due to the absence of PSL (16). Since *pel* is equally up-regulated in the Δ*rsmA* Δ*pel* Δ*psl* strain, demonstrating that the *pel* transcriptional phenotype of Δ*rsmA* is PEL-, PSL-, and c-di-GMP-independent *$p = 0.0002$; **$p = 0.0013$

B. Real-time quantitative PCR at different time points after rifampicin treatment show indistinguishable stabilities of the *pelA* transcript in the WT and Δ*rsmA* strains.

**Fig. S2 RsmA indirectly up-regulates *fleQ* transcript levels.**

A. Single-copy transcriptional *lacZ* fusion constructs reveal down-regulation of *fleQ* transcripts in the Δ*rsmA* background as compared to WT.

B. Real-time quantitative PCR at different time points after rifampicin treatment shows indistinguishable stabilities of the *fleQ* transcript in the WT and Δ*rsmA* strains.

**Fig. S3 EMSA analysis of Hfq binding to *vfr* RNA**

A. *P. aeruginosa* Hfq and *P. putida* Hfq bind identically to *P. aeruginosa vfr* mRNA. Band shifts are indicated by the open arrow heads 1, 2, and 3 as under Fig. 3C. Molar ratios of hexameric Hfq over RNA are indicated; U = unbound RNA

B. Mutant version of *P. putida* Hfq in the distal binding site ($Hfq_{Y25D}$) lack the first band-shift, but the second shift is present. Conversely, the proximal site mutant version of *P. putida* Hfq ($Hfq_{K56A}$) produces the first band but not the second. Neither mutant Hfq proteins produce the third band, which is interpreted to result from two Hfq hexamers simultaneously binding to the A-rich site (Hfq-site 1) and U-rich site (Hfq-site 2) per *vfr* mRNA (Fig. 3A).

**Fig. S4 $P_{pel}$-, $P_{fleQ}$ -and $P_{vfr}$-*lacZ* transcriptional fusion constructs**

A. Simplified plasmid map of mini-CTX *lacZ* as published in (12). The expanded multiple cloning site (MCS) highlights the two restriction sites in blue (EcoRI and BamHI) into which transcriptional fusion fragments (panels B-D) were cloned.

B-D. Intergenic regions that were cloned to produce the transcriptional fusion constructs are shown for $P_{pel}$, $P_{fleQ}$, and $P_{vfr}$ respectively. The regions between the vertical dashed lines were amplified

using the primers (solid black arrows; primer names from Table S2 shown beneath the arrows). The hooked arrows represent the transcriptional start sites and the transcriptional directions of corresponding downstream genes as determined in previous studies (17-19). The specific chromosomal co-ordinate positions are indicated with the numbers corresponding to the *P. aeruginosa* PAO1 genome sequence annotated in www.pseudomonas.com (14, 15).
Items in Fig. S4 are not drawn to scale.

**Table S1**

Bacterial strains and plasmids used in this study.

**Table S2**

Oligonucleotide primers used in this study. Engineered restriction sites are underlined. Genome co-ordinates are based on the annotation of PAO1 on www.pseudomonas.com (14, 15).
